# Host adaptation in gut Firmicutes is associated with sporulation loss and altered colonisation patterns

**DOI:** 10.1101/2020.09.09.289504

**Authors:** Hilary P. Browne, Alexandre Almeida, Nitin Kumar, Kevin Vervier, Anne T. Adoum, Elisa Viciani, Nicholas JR Dawson, Samuel C. Forster, Claire Cormie, David Goulding, Trevor D. Lawley

## Abstract

Human-to-human transmission of symbiotic, anaerobic bacteria is a fundamental evolutionary adaptation essential for membership of the human gut microbiota. However, despite its importance, the genomic and biological adaptations underpinning symbiont transmission remain poorly understood. Here, we show that sporulation ability, which promotes transmission of anaerobic bacteria, has been independently lost in many distinct evolutionary lineages of gut bacteria belonging to the Firmicutes phyla. Analysis of 1358 genome-sequenced Firmicutes reveals loss of sporulation is associated with features of host-adaptation such as genome reduction and specialized metabolic capabilities. Consistent with these data, analysis of 28,000 gut metagenomes from people around the world demonstrates that bacteria now incapable of sporulation are more abundant but less prevalent in the human population compared to spore-forming bacteria. Our results suggest host adaptation in gut Firmicutes is an evolutionary trade-off between transmission range and colonisation abundance, leading to distinct transmission cycles. We reveal host transmission as an underappreciated process that shapes the evolution, assembly and functions of gut Firmicutes.

---

The human gut is colonised by highly adapted bacteria, primarily from the Firmicutes, Bacteroidetes, Actinobacteria and Proteobacteria phyla, that are linked with human health and development^1,2^. Co-evolution between humans and these symbiotic, anaerobic bacteria require that individual bacterial taxa faithfully and efficiently transmit and colonise, as an inability to do either leads to extinction from the indigenous microbiota^3-6^. Key adaptations of symbiotic bacteria in human populations therefore require coordination of colonization and transmission functions. Gut bacteria must be able to colonise above a certain abundance to achieve sufficient shedding levels to ensure onward transmission, and survive in the environment long enough to encounter a susceptible host^5^. Once ingested, gut bacteria must transit through the gastrointestinal tract, contend with the human immune system and compete with indigenous bacteria for nutrients and replicative niches to colonize^5,7^. Transmission of the intestinal microbiota is an ongoing lifelong process starting with maternal transmission around birth and continuing throughout life, especially between co-habiting individuals in regular contact^8-18^. In fact, gut symbiont transmission during co-habitation has a stronger effect on the composition of an individual’s gut microbiota than human genetics^12^, highlighting the importance of transmission in shaping an individual’s microbiota composition and functions. Thus, the transmission cycle of gut bacteria is underpinned by deep evolutionary selection that remains poorly understood.

Spores are metabolically dormant and highly resistant structures produced by Firmicutes bacteria that enhance survival in adverse conditions^19-21^. Sporulation is utilised by anaerobic enteric pathogens such as *Clostridioides difficile* (formerly *Clostridium difficile*) to promote transmission by maintaining environmental viability. Upon ingestion by a new host, the spores germinate in response to intestinal bile acids^22^. We recently demonstrated that at least 50% of the commensal intestinal microbiota also produce resistant spores that germinate in response to bile acids^23^. Hence, the production of spores enhances environmental survival promoting host-to-host transmission and colonisation for a large proportion of the intestinal microbiota^8,23-25^.

Sporulation is an expensive developmental process, dependent on hundreds of genes and takes hours to complete, eventually resulting in the destruction of the original mother cell^21,26,27^. As sporulation is integral to transmission of many gut Firmicutes, we hypothesised that phenotypic loss would confer an advantage linked to an altered transmission cycle no longer reliant on environmental persistence. Loss of sporulation has been demonstrated in experimental conditions under relaxed nutrient selection pressures, indicating maintenance of the phenotype as long as it is beneficial^28,29^. Here, combining large-scale genomic analysis with phenotypic validation of human gut bacteria from the Firmicutes phylum we show that sporulation loss is associated with signature of host adaptation such as genome reduction and more specialised metabolic capabilities. Populations-level metagenomic analysis reveals bacteria no longer capable of sporulation are more abundant but less prevalent compared to spore-formers, suggesting increased colonisation capacity and reduced transmission range are linked to host adaptation within the human intestinal microbiota.

## Results

### Prediction of sporulation capability in gut Firmicutes

We collated 1358 non-redundant, whole genome sequences of Firmicutes bacteria and 72 Actinobacteria, Bacteroidetes and Proteobacteria bacteria for comparative purposes derived from different human different body sites and other environments **(Extended Data Fig. 1, Supplementary Table 1)**. Using this collection, we determined their sporulation capacity, by establishing the presence of 66 sporulation-predictive genes identified using a previously developed machine learning model based on analysis of nearly 700,000 genes and 234 genomes from bacteria with an ethanol sensitive or ethanol resistant phenotype^23^. Genes in this sporulation signature include characterised sporulation-associated genes, characterised genes not previously associated with sporulation and uncharacterised genes that have subsequently been demonstrated to be sporulation-associated^30^.

Interestingly, the largest human gut-associated bacterial families (*Erysipelotrichaceae, Peptostreptococcaceae, Clostridiaceae, Ruminococcaceae* and *Lachnospiraceae*) all display a bimodal sporulation signature score pattern **(Fig. 1a)**. In contrast, other families such as *Lactobacillaceae, Enterococcaceae* and bacteria from the Bacteroidetes, Actinobacteria and Proteobacteria phyla which do not make spores, display only low sporulation signature scores (P<0.0001, Mann-Whitney) **(Fig. 1a, Extended Fig. 2, Supplementary Table 1)**. As bacterial sporulation is believed to have evolved once, early in Firmicutes evolution^19,31^, we classify low scoring sporulation signature genomes (average sporulation signature score of genomes in family <50%) as Former-Spore-Formers (FSF) (i.e. Firmicutes that have lost the capability to produce spores) and high scoring sporulation signature genomes (average sporulation signature score of genomes in family ≥50%) as Spore-Formers (SF). This classification is a refinement of our previously established sporulation signature that accounts for the different sporulation machinery between taxonomically different bacterial families^23^.

**Figure 1:**
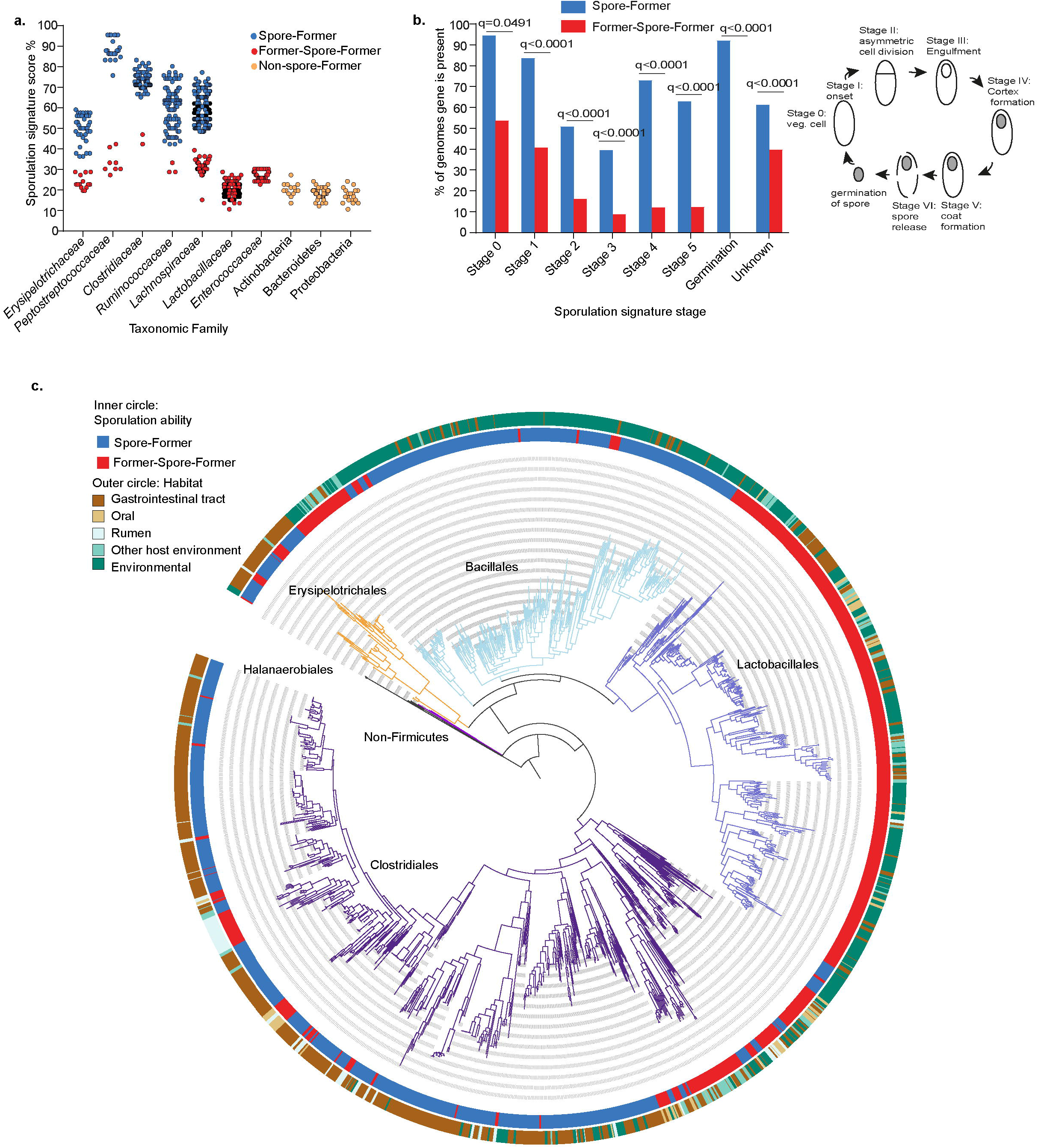
Sporulation loss in distinct evolutionary lineages of gut Firmicutes. **(a)** Prediction of sporulation capability in human-associated Firmicutes families based on the presence of 66 sporulation associated genes. *Erysipelotrichaceae, Peptostreptococcaceae, Clostridiaceae, Ruminococcaceae* and *Lachnospiraceae* families have a bimodal pattern with genomes either having a high scoring (average sporulation signature score of genomes in family ≥50%) (classified as Spore-Formers) or a low scoring sporulation signature score (average sporulation signature score of genomes in family <50%) (classified as Former-Spore-Formers). *Lactobacillaceae, Enterococcaceae*, Actinobacteria, *Bacteroidetes* and Proteobacteria are all non-spore-forming and contain low scoring genomes. **(b)** Presence and absence of sporulation signature genes was determined in gut-associated Spore-Formers (SF) (n=456), and Former-Spore-Formers (FSF) (n=118) bacteria. FSF have less sporulation signature genes for all sporulation stages compared to SF (all stages q< 0.0001, except for stage 0 q=0.0491, Fishers exact test, adjusted for multiple testing). Cartoon describes sporulation stages. **(c)** Phylogeny of the Firmicutes constructed from 40 universal protein coding genes extracted from 1358 whole genome sequences. Sporulation has been lost at large taxonomic scales, (*Lactobacillales* order) and at small taxonomic scales (within host-associated *Erysipelotrichales* and *Clostridiales* orders). Major taxonomic orders are indicated by branch colours and name, black branches at base of phylogeny represent non-Firmicutes root derived from Actinobacteria genomes.

To investigate the genetic processes and selective forces underlying sporulation loss in human gut symbionts, we combined genomes from gut-associated SF and FSF bacteria (SF n=455, FSF n=118), determined the presence of the 66 sporulation signature genes and assigned them to their respective sporulation stage. As expected, FSF genomes contain fewer sporulation signature genes for each sporulation stage compared to SF genomes (all stages, q< 0.0001 except for Stage 0 (q= 0.0491, Fishers exact test) **(Fig 1b)**. Early-stage (Stage 0 and I) sporulation genes which are unknown in function or have pleiotropic, non-sporulation related functions are maintained to a greater degree compared to later-stage sporulation genes in FSF genomes (Stage 0 sporulation genes are, on average, present in 53.7% of FSF genomes, Stage I present in 40.7% of FSF genomes). Later-stage sporulation signature genes that are sporulation-specific are absent to a greater degree in FSF genomes (Stage II sporulation genes are, on average, present in 16.2% of FSF genomes, Stage III genes present in 8.8% of FSF genomes, Stage IV genes are present in 12% of FSF genomes, Stage V genes are present in 12.2% of FSF genomes and germination stage genes are absent from FSF genomes). Hence, sporulation-specific genes may be lost as there is no advantage in maintaining them.

We next phenotypically validated the lack of sporulation in gut-associated FSF. We exposed cultures of 41 phylogenetically diverse species from 6 different Firmicutes families, SF (n=26) and FSF (n= 15) to 70% ethanol for 4 hours and then cultured on YCFA nutrient media with sodium taurocholate to stimulate germination of ethanol-resistant spores^23,32^ **(Supplementary Table 1)**. In addition, to account for bacteria that require intestinal signals to produce spores not present in our experimental conditions, we also recorded whether these species were originally cultured from ethanol-exposed faecal samples^23^. Only SF species (12 out of 26) were successfully cultured after ethanol exposure. A further 9 were not cultured after ethanol exposure but were originally isolated from ethanol exposed faeces, highlighting that for some species sporulation is not induced *in vitro*. Taken together, 21 of 26 total (81%) produce ethanol resistant spores **(Extended Data Fig. 3a)**. No FSF survived ethanol exposure (0/15 (0%)) and none were originally isolated from ethanol exposed faeces **(Supplementary Table 1**). Transmission Electron Microscopy (TEM) imaging confirmed the presence of spores in spore-forming bacteria only **(Extended Data Fig. 3b)**, including those with a sporulation signature score below the previously used threshold of 50%^23^. Thus, we demonstrate loss of sporulation-specific genes leading to an absence of spores in distinct evolutionary lineages of bacteria, creating Former-Spore-Formers.

### Large-and small-scale phylogenetic loss of sporulation in Firmicutes

The differences in sporulation gene content within and between families indicates a divergence in sporulation capacity between distinct lineages, raising interesting questions regarding the phylogenetic and evolutionary relationship between sporulating and non-sporulating bacteria. We next generated a core gene phylogeny, of the 1358 Firmicutes genomes, and mapped sporulation capability to better understand the evolution of sporulation in human gut symbionts **(Fig. 1c)**. Our analysis places the non-gut-associated, SF order Halanaerobiales at the base of the phylogeny^33^. Subsequent, large-scale loss of sporulation within taxonomic orders such as the *Lactobacillales* is evident (all 344 genomes are predicted to be FSF), which has been observed before and attributed to adaptation to nutrient-rich habitats^19,34^. Interestingly, we also observe smaller scale sporulation loss within multiple distinct clades of the host-associated *Erysipelotrichaceae*, (26% are FSF), *Peptostreptococcaceae* (26% are FSF) and *Lachnospiraceae* (18% are FSF) families^35^. Within host-associated bacteria, sporulation has been lost to a greater degree in non-gut habitats (96.6% of oral-associated bacteria are FSF, 84.1% of rumen-associated bacteria are FSF, while only 20.7% of gut-associated bacteria are FSF). Thus, although sporulation is a core function of the human gut microbiota, as it enhances faecal-oral transmission, we reveal loss of sporulation capability in multiple distinct lineages of human-associated Firmicutes bacteria.

### Genome reduction in gut Former-Spore-Formers

Genome reduction is a feature of host-adaptation and is characterised by a loss of genes not required to survive in an ecosystem^36-39^. To determine if loss of sporulation genes in FSF is associated with broader genome decay, we next compared genome sizes of gut FSF and SF bacteria. FSF (n=118) have, on average, genomes that are 36% smaller than SF bacteria (n=456) (P< 0.0001, Mann-Whitney) **(Fig. 2a)**. The same trend is present in FSF genomes in *Erysipelotrichaceae* (38.9% smaller, P<0.0001, Mann-Whitney), *Peptostreptococcaceae* (40.1% smaller, P=0.0002, Mann-Whitney) *and Lachnospiraceae* families (15.6% smaller, P<0.0001, Mann-Whitney) which contain both SF and FSF bacteria **(Extended Data Fig. 4a)**. A low genetic redundancy is another feature of host adaptation and is associated with occupation of stable or constant niches within an ecosystem^39^. Within the same three families, FSF have a lower percentage of paralogous genes in their genomes in comparison to SF bacteria (*Erysipelotrichaceae* P<0.0001, *Peptostreptococcaceae* P=0.0002, and *Lachnopsiraceae* P< 0.0001, Mann-Whitney) **(Extended Data Fig. 4b)**. Thus, within FSF bacteria, loss of sporulation genes is associated with broader genome decay, evidence for host adaption and an altered transmission cycle.

**Figure 2:**
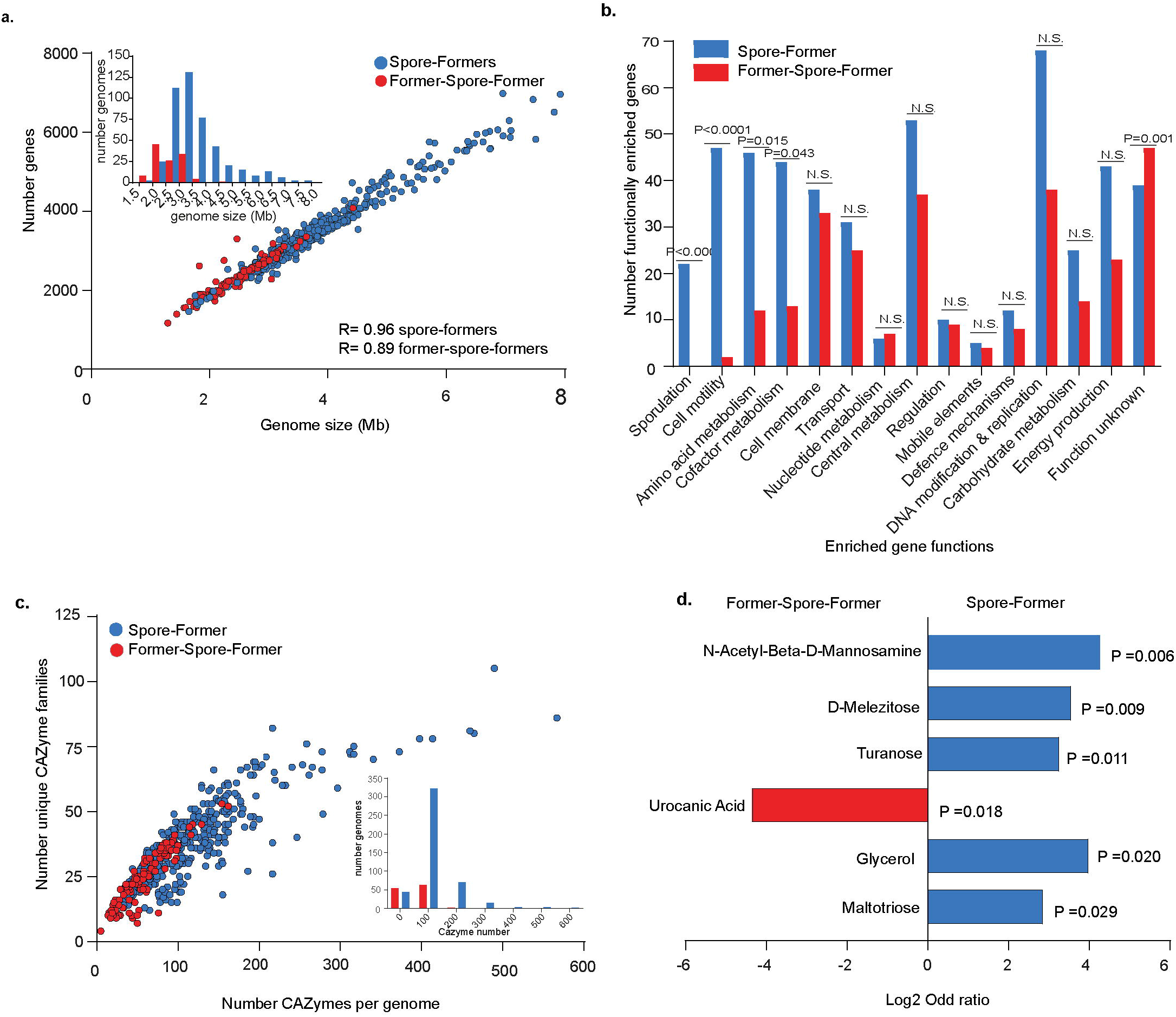
Genome reduction and metabolic specialization during host-adaptation by gut Former-Spore-Formers. **(a)** A marker of host adaptation, genomes of gut FSF (n=118) are smaller than SF genomes (n=456) (P< 0.0001, Mann-Whitney test), with a strong correlation between genome size and gene number, Spearman rho, R= 0.96 for SF and R= 0.89 for FSF. Inset shows distribution by genome size. **(b)** Functional enrichment analysis revealed 489 enriched gut SF genes and 272 enriched FSF genes. Enriched genes were grouped by functional classes. Graph presents comparison of enriched gene numbers in their functional class and ordered by decreasing statistical significance. Motility, amino acid and cofactor metabolism and sporulation functional classes are statistically more enriched in gut Spore-Formers (SF) compared to Former-Spore-Formers (FSF). No functional classes are more enriched in FSF, Fisher’s exact test, N.S not significant **(c)** FSF encode a smaller number of total CAZymes and a smaller number of CAZyme families per genomes compared to gut SF (P<0.0001 for both total number and family number, Welch’s t-test). Inset shows distribution by CAZyme number. **(d)** *Erysipelotrichaceae* FSF have a more restricted carbohydrate utilisation profile compared to *Erysipelotrichaceae* SF. The ability of FSF (n=4) and SF (n=4) to use 95 different carbon sources was tested. N-Acetyl-Beta-D-Mannosamine (P=0.006) (a precursor of sialic acid), D-Melezitose (P=0.009), turanose (P=0.011), glycerol (P=0.020) and maltotriose (P=0.029) are metabolised to a statistically significant greater degree by SF whereas urocanic acid (a derivative of histidine) was metabolised to a statistically significant greater degree by FSF (P=0.018), based on Fisher’s exact test).

### Metabolic specialization during host-adaptation by gut Former-Spore-Formers

We next carried out functional enrichment analysis to define genome-wide adaptive features differentiating human gut-associated SF and FSF bacteria. In total 489 genes were enriched in SF and 272 were enriched in FSF. We assigned these genes to functional classes based on their annotation and compared functional classes enriched in both groups **(Fig. 2b, Supplementary Table 2)**. Cell motility (P<0.001), amino acid metabolism (P=0.0148), cofactor metabolism (P=0.043), and sporulation (P<0.001, Fisher’s exact test) functional classes were statistically significantly enriched in SF compared to FSF. No functional class was enriched in FSF compared to SF.

Within SF, we observe a tendency towards biosynthesis of metabolites compared to transport of metabolites in FSF. The majority (31 of 46) of enriched genes with amino acid metabolism functions in SF are biosynthesis-associated (including histidine, methionine, leucine and isoleucine). By comparison, 6 of 12 enriched genes with amino acid metabolism functions in FSF are transport-associated (only 3 are biosynthesis-associated). Similarly, 41 of 44 enriched genes with cofactor metabolism functions in SF are biosynthesis-associated, including cobalamin (Vitamin B12) (n=19 genes of 23 total required for cobalamin biosynthesis) **(Extended Data Fig. 4c)**. Cobalamin is primarily acquired by external transportation by gut bacteria and is required for important microbial metabolic processes, including methionine biosynthesis^40,41^. Species auxotrophic for cobalamin rely on sharing from cobalamin producers, hence these functions in SF may promote stability within the intestinal microbiota by providing essential metabolites^42,43^.

Within FSF, no cobalamin biosynthesis genes are enriched, but 2 (*BtuB* and *BtuE*) are associated with cobalamin transport. Also within the cofactor metabolism class, FSF are enriched in 4 genes associated with Vitamin K2 (menaquinone) biosynthesis. Interestingly, microbial production of cobalamin is unlikely to benefit human hosts due to an inability to absorb it, unlike menaquinone where absorption is possible^40^. Amino acid and cofactor transport may therefore provide an adaptive efficiency for FSF bacteria avoiding the cost of internal biosynthesis. Thus, for certain key metabolites, host adaptation by gut bacteria may be characterized as lifestyle shift from “producer” to “scavenger”, potentially promoting colonization of distinct metabolic niches.

### Restricted carbohydrate metabolism in Former-Spore-Formers

Within SF, 25 functionally enriched genes are annotated with carbohydrate metabolism functions compared to 14 in FSF **(Fig. 2b)**. As carbohydrates are the main energy source for gut bacteria, we next annotated the Carbohydrate-Active Enzymes (CAZymes) within Firmicutes gut bacteria. On average, SF genomes have a larger total number of CAZymes and a larger number of different CAZyme families compared to FSF genomes (total number CAZymes: 112 on average per SF genome compared to 57.51 per FSF genome, number CAZymes families: 37 on average per SF genome compared to 24.16 per FSF genome) (total number and family number of CAZyme P<0.0001, Welch’s t-test) **(Fig. 2c)**. Thus, SF encode a broader repertoire of CAZymes, suggesting a greater saccharolytic ability.

The *Erysipelotrichaceae* are a phylogenetically distinct bacterial family within the *Erysipelotrichales* order that remain poorly characterised despite being both health- and disease-associated in humans^44^. Importantly, in our dataset, the *Erysipelotrichaceae* contain multiple gut-associated SF and FSF specie**s (Extended Data Fig. 5a)**. Reflecting the broader pattern in the Firmicutes **(Fig. 2c)**, *Erysipelotrichaceae* gut SF encode a larger total number and a larger number of CAZyme families compared to *Erysipelotrichaceae* gut FSF **(Extended Data Fig. 5b)** (total number CAZymes: 115 on average per SF genome compared to 57 per FSF genome, number CAZymes families: 34.85 on average per SF genome compared to 23 per FSF genome) (total number and family number of CAZyme P<0.0001 and P=0.0001 respectively, Welch’s t-test). We next chose to use this family to phenotypically validate our genomic analysis results showing a broader carbohydrate metabolic profile in SF. We inoculated phylogenetically diverse bacteria from *Erysipelotrichaceae* SF (n=4) and FSF (n=4)^23,45^ **(Extended Data Fig. 5a, Supplementary Table 3)** in Biolog AN MicroPlates containing 95 different carbon sources. Growth was detected for 78 different carbon sources (59 carbon sources for FSF and 69 carbon sources for SF) **(Supplementary Table 4)**. When clustered into broad carbon source groups, FSF were more limited in their capacity to utilise both carbohydrates (P<0.0001, Fisher’s exact test) and amino acids (P=0.003, Fisher’s exact test) **(Extended Data Fig. 5c)**, consistent with our genomic analysis **(Fig. 2c & Extended Data Fig. 5b)**.

At the individual carbon source level, FSF also had a reduced metabolic capability compared to SF bacteria **(Fig. 2d)**. Urocanic acid, a derivative of histidine, whose metabolism is linked to short chain fatty acid production, was the only carbon source metabolised to a statistically significant greater degree by FSF (P=0.018, Fisher’s exact test), suggesting metabolism of specific amino acids present in the intestinal environment may provide colonisation-associated advantages in the FSF *Erysipelotrichaceae*. Enriched metabolism of carbon sources by SF include glycerol (P=0.020, Fisher’s exact test), which requires cobalamin as a cofactor for its metabolism^46^, and N-Acetyl-Beta-D-Mannosamine (β-ManNAc) (P=0.006, Fisher’s exact test), a derivative of sialic acid. Thus, we provide evidence of a specialised metabolic capability linked to host adaptation in Firmicutes gut bacteria.

### Former-Spore-Formers display increased colonisation abundance in human populations

Taken together, our genotypic and phenotypic results indicate that the broader metabolic and functional capabilities of SF reflect a more generalist lifestyle, compared to the reduced capabilities of FSF which we propose are adapted to a more stable and specialised lifestyle. We next hypothesised that an inability to make spores in FSF bacteria would limit their environmental survivability and, as a result, reduce their transmission range leading to a lower prevalence in human populations compared to SF bacteria. To investigate this, we calculated the prevalence of SF and FSF bacteria in 28,060 faecal metagenomes representing human populations from 6 continents and healthy and disease states **(Supplementary Table 5)** by reference genome-based mapping to our annotated genomes for SF and FSF bacteria^45^. Importantly, we found that FSF were significantly less prevalent (P=0.0098, two-tailed Wilcoxon rank-sum test) compared to SF **(Fig. 3a)**. Interestingly, this trend is also present in the broader intestinal microbiota. Non-Spore-Formers (NSF) (non-Firmicutes that never made spores and FSF) are significantly less prevalent than SF (P=0.0034, two-tailed Wilcoxon rank-sum test) **(Extended Data Fig. 6a)**. These results provide evidence that loss of sporulation limits the transmission range of FSF in human populations.

**Figure 3:**
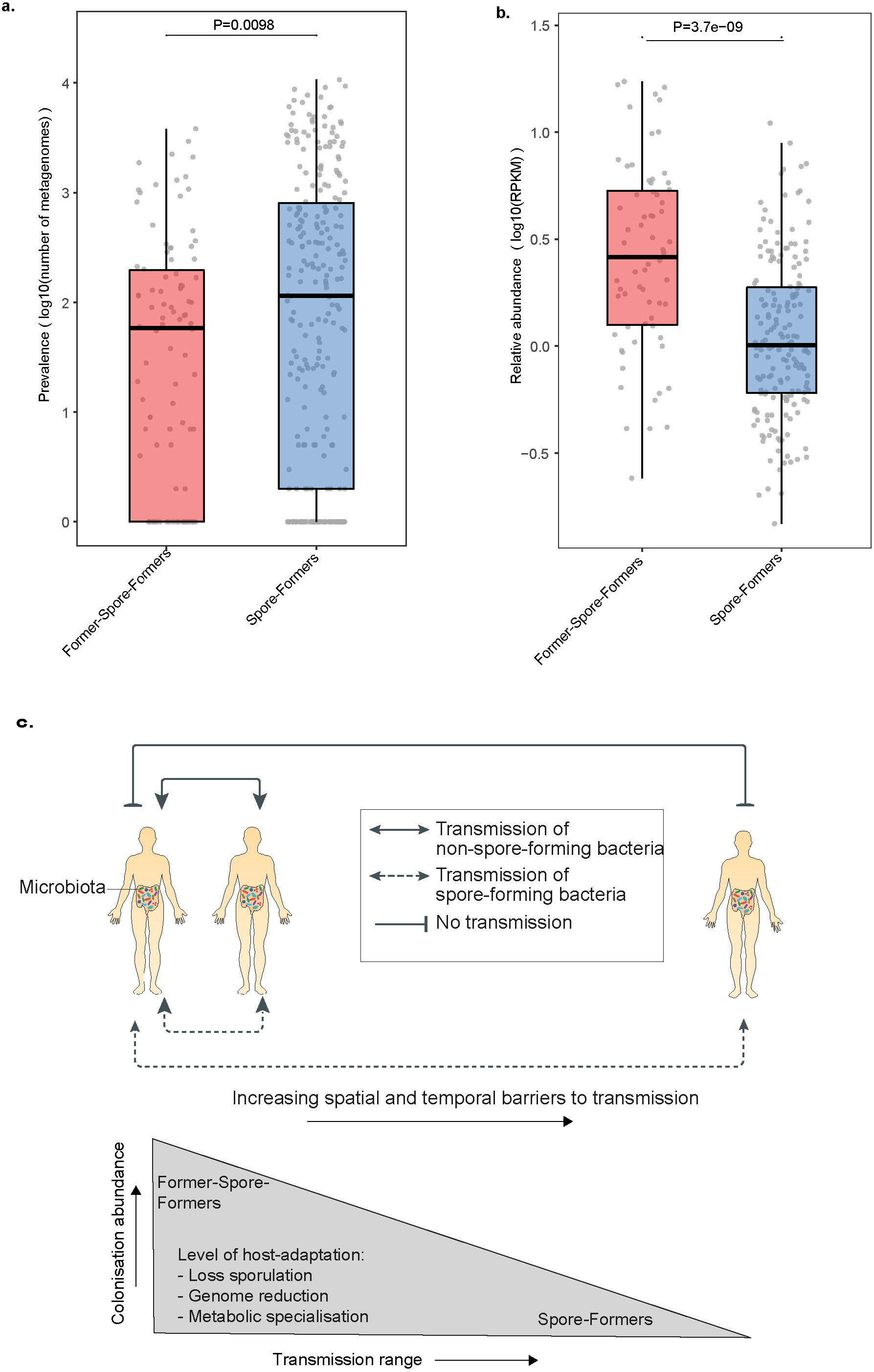
Host adaptation is associated with reduced prevalence and higher colonisation abundance levels in Former-Spore-Formers. **(a)** Former-Spore-Formers are less prevalent (P=0.0098, two-tailed Wilcoxon rank-sum test) compared to Spore-Formers within 28,060 faecal metagenomes reflecting a reduced transmission range. **(b)** While less prevalent, Former-Spore-Formers are more abundant (P<0.001, two-tailed Wilcoxon rank-sum test) compared to Spore-Formers within the same 28,060 faecal metagenomes reflecting greater host adaptation and an ability to colonise to higher levels. Each dot represents an individual species. Box lengths represent the IQR of the data, and the whiskers the lowest and highest values within 1.5 times the IQR from the first and third quartiles, respectively. **(c)** Host adaptation is associated with an altered transmission cycle in gut Firmicutes. Former-Spore-Formers (FSF) are more host-adapted compared to Spore-Forming (SF) bacteria as measured by their reduced genome size and genomic redundancy and their more specialist metabolic capabilities. This greater level of host-adaptation corresponds to an ability to colonise to higher abundance levels which promotes onward direct transmission to hosts in close proximity, By contrast, SF are less host-adapted and colonise to lower abundance levels. SF transmission cycle relies on the production of resilient spores that promote environmental persistence, thus they are capable of colonising a greater proportion of hosts and have a greater prevalence in human populations. Figure adapted from^5^.

Our genotypic and phenotypic results indicate that FSF may be more specialized in metabolism and therefore may have a growth advantage in the human gut. We next examined if host adaptation in FSF correlates with an ability to colonise humans to higher levels than SF bacteria. Using reference genome-based mapping, we detected FSF at a significantly higher relative abundance than SF in the human intestinal microbiome (P<0.001, two-tailed Wilcoxon rank-sum test) **(Fig. 3b)**. We also observe the same trend when expanding this analysis to the entire intestinal microbiota; NSF are significantly more abundant than SF **(Extended Data Fig. 6b)** (P<0.001, two-tailed Wilcoxon rank-sum test). A higher abundance of FSF would promote onward transmission to hosts in close proximity and over short periods, by increasing the levels of excreted bacteria, promoting bacteria maintenance in the local human population. Hence, an absence of sporulation in FSF is correlated with higher abundance levels in the intestinal microbiota, potentially reflecting distinct transmission and colonisation strategies in the intestinal microbiota between bacteria that are capable and incapable of sporulation.

## Discussion

Here, we show evidence that gut Firmicutes commonly lose their ability to make spores during host adaptation. FSF bacteria are less resilient compared to spore-formers, which limits environmental survival, resulting in an altered transmission cycle^5,23^. Within closely interacting, social groups of baboons, non-spore-forming, anaerobic bacteria (including FSF) are shared to a greater degree than spore-forming bacteria^47^, suggesting a transmission cycle that relies on close contact between hosts to limit bacterial exposure to adverse environmental conditions. Furthermore, colonisation to high abundance levels which is a feature of Former-Spore-Formers (and non-spore-forming bacteria within the broader intestinal microbiota) will promote transmission by ensuring high shedding levels in faecal matter that would increase the chances of a successful host colonisation event^48^. Indeed, a greater incidence of transmission between mother and infants is observed for FSF compared to SF^17^. Thus, the transmission cycle of FSF bacteria is highly evolved, potentially relying on high level colonisation abundance to facilitate transmission over short distances and time frames **(Fig. 3c)**.

By contrast, environmental persistence of resilient spores removes the need for direct transmission between hosts in close contact. The larger genomes of SF, encoding a broader metabolic capability, also indicate a more generalist lifestyle adept at surviving in different hosts and environments^49^. Previous studies have shown that human acquisition of SF in early life occurs to a greater degree from environmental sources compared to non-spore-forming bacteria which are maternally-acquired^8,17,50^. Thus, the transmission cycle of SF relies on the production of resilient spores which increases the proportion of individuals that can potentially be colonised and which is reflected in the greater prevalence of SF in human populations. Hence, SF have a larger transmission range compared to non-spore-forming bacteria **(Fig. 3c)**.

We propose there is an evolutionary trade-off between high level colonisation abundance mediated by host adaptation and transmission range promoted by sporulation within gut Firmicutes. Sporulation is an expensive biological process requiring synchronisation of hundreds of genes, hence it is likely to be lost if no longer needed^28^. Lowly abundant SF bacteria are more at risk of extinction or expulsion from the intestinal microbiota, therefore sporulation may be maintained as it increases chances of survival once expelled^51^. Alternatively, loss of sporulation may lead to a different evolutionary trajectory centred on host adaptation, high level colonisation abundance and a more specialised transmission cycle. Hence, we reveal different levels of host adaptation exist within gut Firmicutes impacting transmission and colonisation processes.

---

**Extended Data Figure 1:**
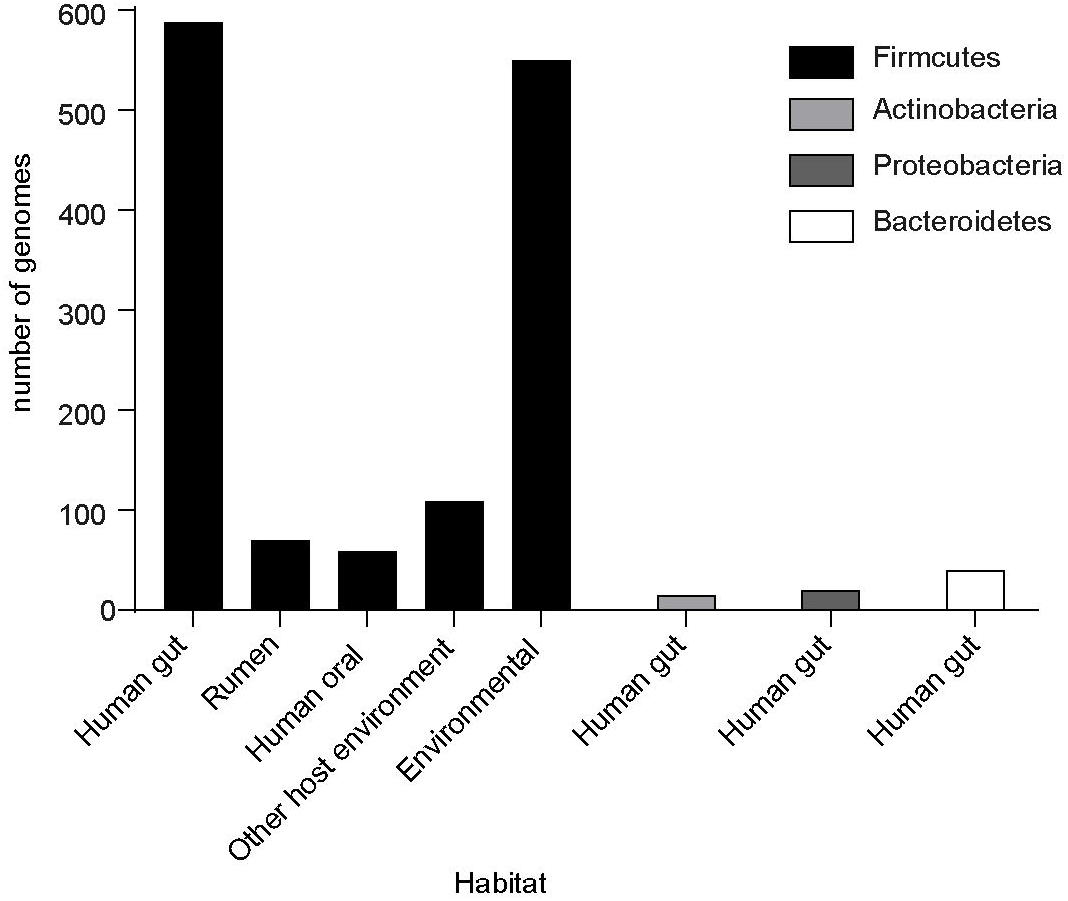
Environmental distribution of genomes from Firmicutes bacteria. Firmicutes genomes (n=1358) are derived from bacteria present in a diverse range of host and non-host-associated habitats. Actinobacteria (n=14), Proteobacteria (n=19) and Bacteroidetes (n=39) genomes, all derived from gut bacteria, were included in analysis for comparative purposes.

**Extended Data Figure 2:**
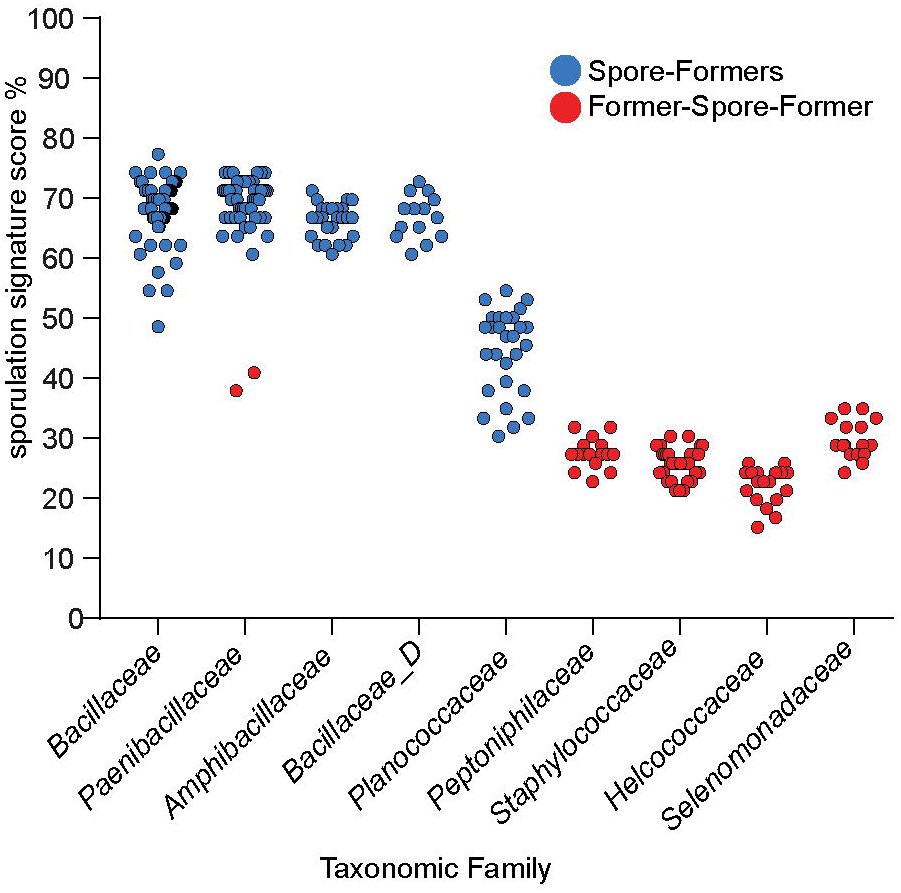
Prediction of sporulation capability in Firmicutes. **(a)** Prediction of sporulation capability in Firmicutes families based on the presence of 66 sporulation associated genes. These families have either a high sporulation signature score (average sporulation signature score of genomes in family ≥50%) (classified as Spore-Formers), a low sporulation signature score (average sporulation signature score of genomes in family <50%) (classified as Former-Spore-Formers) or a bimodal pattern with both high and low scoring sporulation signature genomes.

**Extended Data Figure 3:**
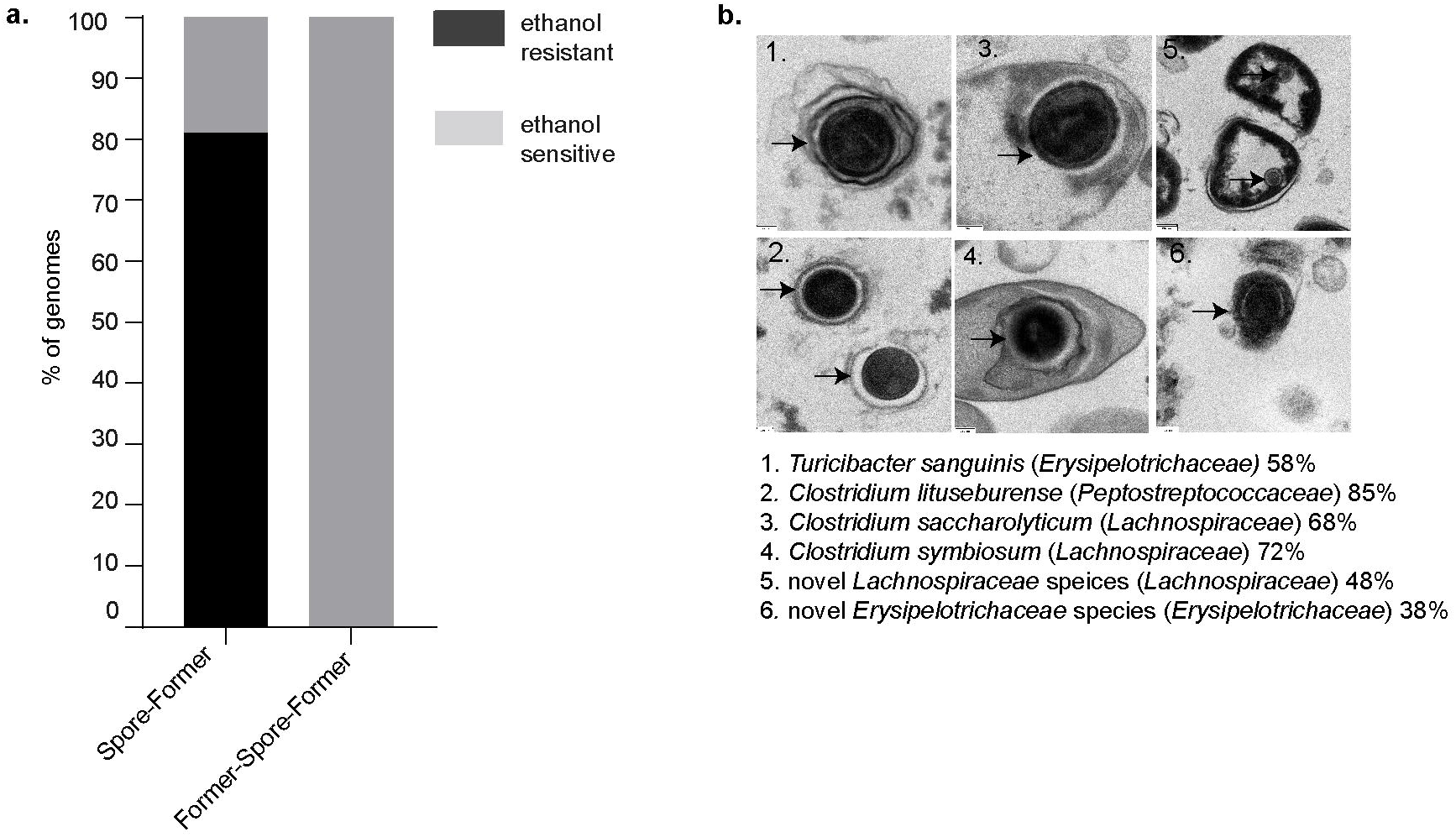
Phenotypic validation of sporulation capability predictions. Phenotypic validation of loss of sporulation using exposure of isolates to ethanol and isolation from ethanol treated faeces **(a)** and Transmission Electron Microscopy (TEM) **(b)**. Spores were only identified in bacteria predicted to be spore-forming. Arrows indicates spores. Species name corresponding to TEM image is indicated along with corresponding family in brackets and sporulation signature score.

**Extended Data Figure 4:**
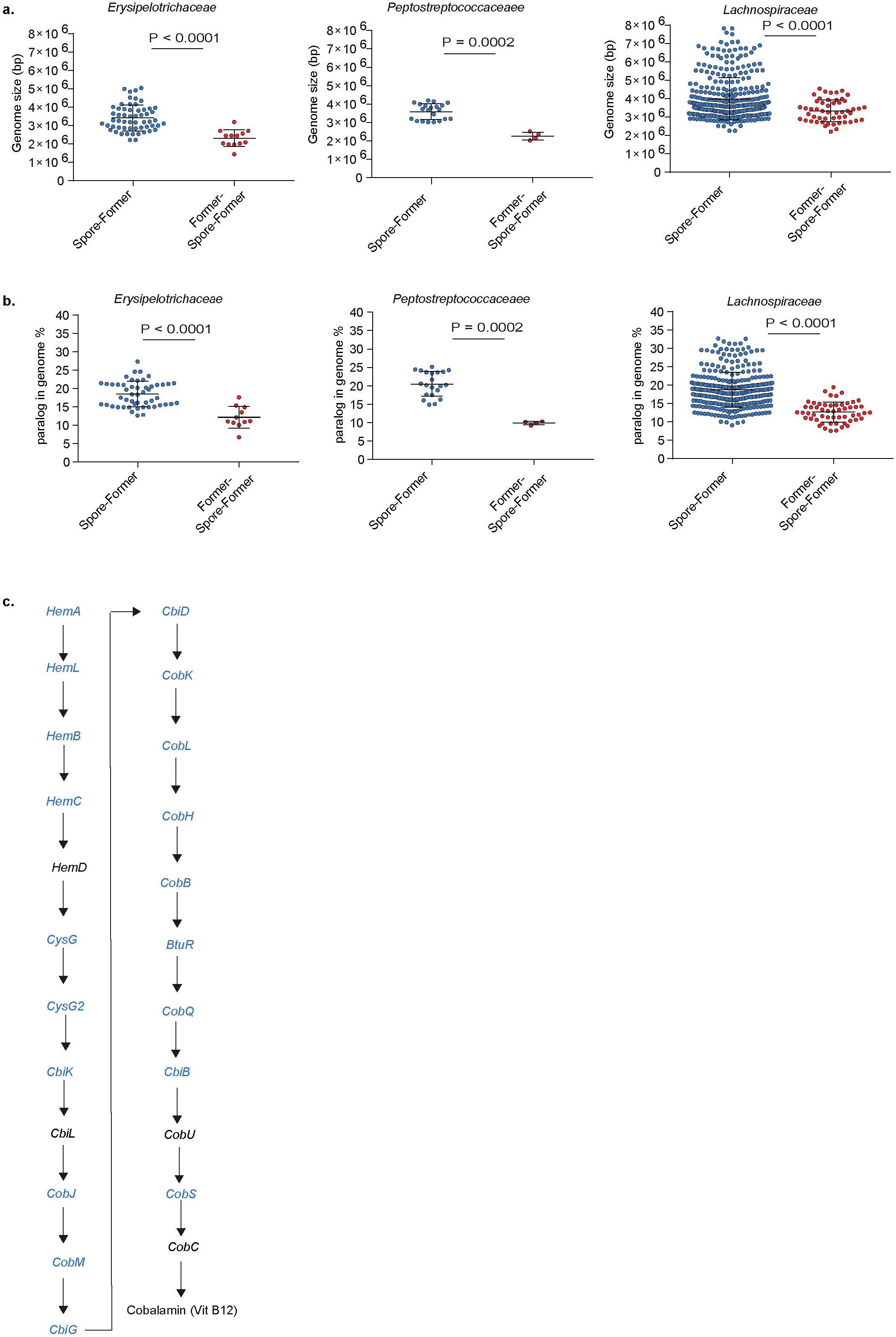
Genome reduction and metabolic specialization during host-adaptation by gut Firmicutes. Former-Spore-Formers (FSF) of host-associated bacterial families, *Erysipelotrichaceae, Peptostreptococcaceae* and *Lachnospiraceae* have more reduced genomes compared to Spore-Formers (SF) within the same family as measured by a smaller genome size **(a)** (*Erysipelotrichaceae* SF gut vs. FSF gut P< 0.0001, *Peptostreptococcaceae* SF gut vs. FSF oral P=0.0002, *Lachnopsiraceae* SF gut vs. FSF oral and rumen P< 0.0001, Mann-Whitney), mean with SD, and by encoding less paralogs **(b)** (*Erysipelotrichaceae* SF gut vs. FSF gut P<0.0001, *Peptostreptococcaceae* SF gut vs. FSF oral P=0.0002, *Lachnopsiraceae* SF gut vs. FSF oral and rumen P< 0.0001, Mann-Whitney), mean with SD. **(c)** SF are enriched in 19 of 23 genes (marked in blue) directly required for cobalamin (Vitamin B12) biosynthesis unlike FSF which are not enriched for cobalamin biosynthesis genes.

**Extended Data Figure 5:**
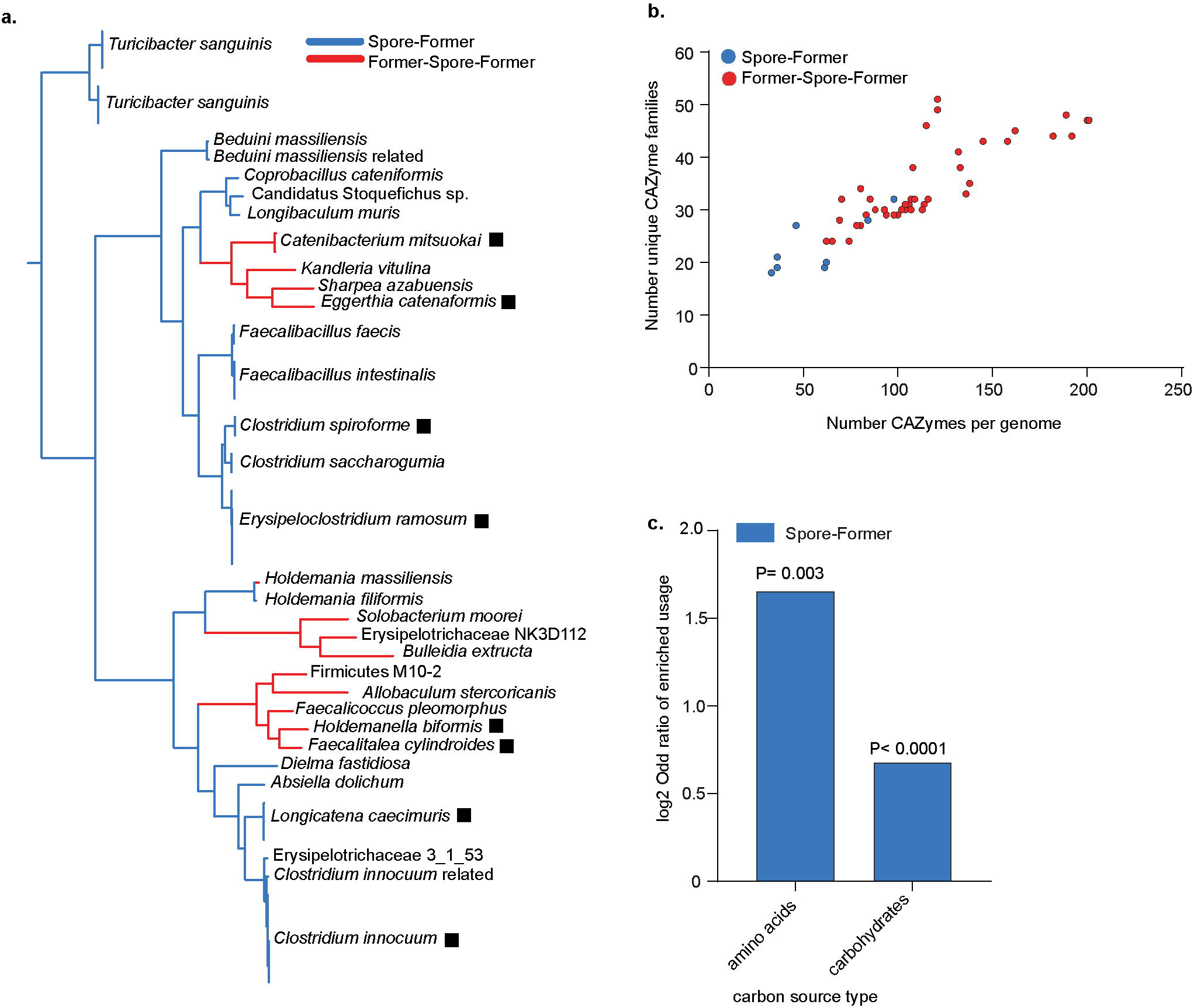
*Erysipelotrichaceae* Former-Spore-Formers have a reduced carbohydrate metabolism profile compared to *Erysipelotrichaceae* Spore-Formers. **(a**) Phylogeny of the *Erysipelotrichaceae* family, the 32 distinct species names are indicated. Species whose name is followed by a rectangle were selected for phenotypic analysis. Novel species are denoted by closest relative followed by term ‘related’. **(b)** *Erysipelotrichaceae* gut FSF encode a smaller number of total CAZymes and a smaller number of CAZyme families per genomes compared to *Erysipelotrichaceae* gut SF (P<0.0001 and P=0.0001 for total number and family number of CAZymes respectively, Welch’s t-test). **(c)** The ability of FSF (n=4) and SF (n=4) to use 95 different carbon sources was tested. Carbon sources were grouped by category and the log2 odds ratio was calculated. Carbohydrates (45 in total) (P<0.0001) and amino acids (17 in total) (P=0.003, Fishers exact test) were utilised less by *Erysipelotrichaceae* FSF compared to SF. No carbon source group were utilised more by FSF compared to SF.

**Extended Data Figure 6:**
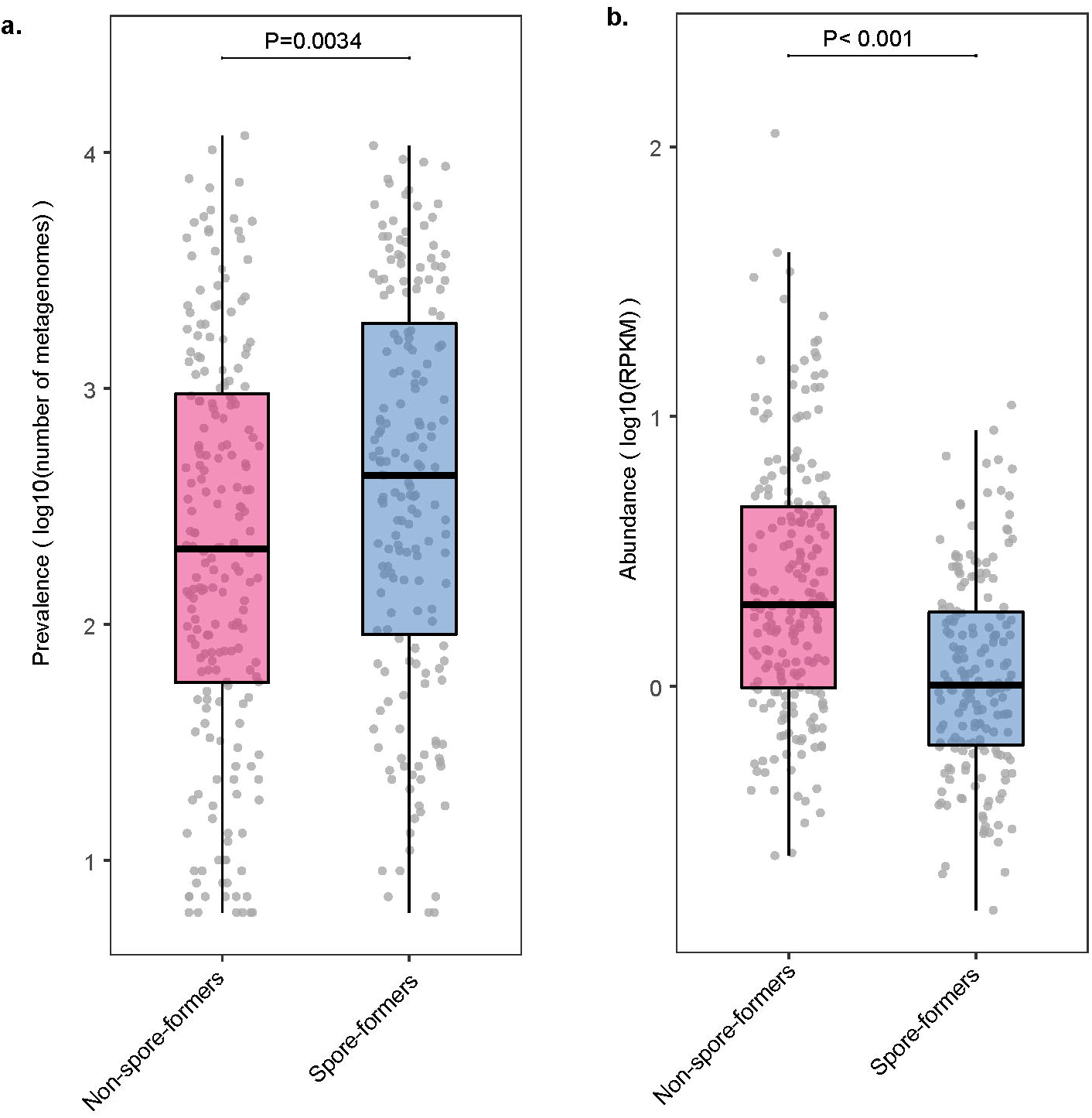
Gut bacteria incapable of sporulation are less prevalent but more abundant than gut spore-forming bacteria. **(a)** Intestinal non-spore-forming bacteria (NSF) (FSF Firmicutes and non-Firmicutes that never made spores such as Bacteroidetes, Actinobacteria and Proteobacteria bacteria) are less prevalent than spore-forming bacteria (SF) in human populations in 28,060 metagenomes. **(b)** While less prevalent, intestinal NSF bacteria are more abundant within the same 28,060 metagenomes. Each dot represents an individual species. Box lengths represent the IQR of the data, and the whiskers the lowest and highest values within 1.5 times the IQR from the first and third quartiles, respectively.

## Methods

### Determination of sporulation ability

We previously identified 66 genes that are predictive for the formation of ethanol-resistant spores using a machine learning approach based on comparison of genomes derived from 234 bacteria with an ethanol resistant or ethanol sensitive phenotype^23^. We applied a strict minimum cut-off of 50% in sporulation signature score (i.e. at least 33 of the 66 sporulation signature score genes present) to classify a genome as capable of sporulation. Here, using a larger data-set of Firmicutes from different environments (not just the gut) we applied the sporulation signature score, assessing sporulation capability on a taxonomic family basis. Families were defined using the GTDB database^52^ and phylogenetic placement in **Fig. 1c**. The presence of the 66 sporulation associated genes were then determined using tblastn of the genome sequences against the amino acid sequence of the 66 sporulation signature genes (e-value 1e-05 and 30% identity). Genomes in families clustered together with either high sporulation signature score scores (classified as spore-forming if average sporulation signature score of genomes in family is ≥ 50%), low sporulation signature scores (classified as non-spore-forming if average signature score of genomes in family is < 50%) or a bimodal pattern with both high (classified as spore-forming) and low scoring signature score (classified as non-spore-forming). This classification system is less stringent than the previously used cut-off of 50% and is supported by the generation of TEM images **(Extended Data Fig. 3b)**, showing spores for genomes of bacteria with a sporulation signature score less than 50% and by literature searches. The average of the sporulation signature score for the *Planococcaceae* family is 45% but literature searches identified this family as spore-forming and all *Planococcaceae* genomes contain *spo0A*, the master regulator gene of sporulation. Hence, it was classified as spore-forming. Furthermore, as all bacteria in the *Planococcaceae* in this dataset are found in non-host environments, it was not a part of the main analysis of gut genomes.

To determine loss of sporulation genes in genomes, the amino acid sequence of the 66 sporulation signature proteins was blasted against whole genome sequences using tblastn, (1e-05, minimum identity 30%). The sporulation signature genes were assigned to specific sporulation stages if known and the percentage of genomes containing genes from each sporulation stage was calculated.

### Phylogenetic analysis

1687 Firmicutes genomes from the NCBI curated RefSeq database, in addition to whole genome sequences from intestinal isolates from the Human Microbiome Project, a comprehensive study describing the first 1000 intestinal cultured species of the intestinal microbiota and an in-house collection of whole genome sequences derived from our intestinal bacterial culture collection were used^23,45,53-55^. Genomes were annotated using the pipeline described in Page *et al*.^56^. Redundant genomes were removed and CheckM^57^ was then used to filter genomes with less than 90% completeness and greater than 5% contamination leaving 1358 genomes for phylogenetic analysis. Actinobacteria genomes (n=5) from our culture collection were used to root the tree-*Collinsella aerofaciens* (GCA_001406575.1), *Bifidobacterium adolescentis* (GCA_001406735.1), novel *Collinsella* species (GCA_900066465.1), *Collinsella aerofaciens* (GCA_001404695.1) and *B. pseudocatenulatum* (GCA_001405035.1). The fetchMG program^55^ was used to extract 40 universal genes from the genomes. The resulting amino acid sequences were aligned using mafft (v7.205)^58^, and gaps representing poorly aligned sequence were removed using the Gblocks script (v0.91b)^59^ leaving an alignment 6048 amino acids in length. A maximum likelihood phylogeny was constructed using FastTree^60^ (version 2.1.9) using the Jones-Tayler-Thorton (JTT) model of amino-acid evolution and 20 rate categories per site. All bootstrap values using the Shimodaira-Hasegawa test, to at least the family level of the phylogeny are greater than 0.7. This phylogenetic structure is congruent with other large phylogenies such as that implemented in AnnoTree^61^ and derived from GTDB^52^. All phylogenies were viewed using iTOL^62^. Habitat origins of isolate genomes were determined using literature searches and available information on NCBI. The *Erysipelotrichaceae* phylogeny was extracted from the main Firmicutes phylogeny.

### Ethanol shock test

Species for the ethanol shock test to validate spore-formation characterisation were selected based on phylogenetic diversity (6 different families were tested, *Enterococcaceae, Streptococcaceae, Lactobacillaceae, Erysipelotrichaceae, Lachnospiraceae* and *Peptostreptococcaceae*) and a wide range in sporulation signature scores (36% to 95% for SF and 15% to 29% for FSF). Isolates were streaked from frozen glycerol stocks and then grown overnight in 10ml broth containing YCFA media, a nutrient growth media formulated to culture gut bacteria^32^. Culturing took place in anaerobic conditions in an A95 Whitley Workstation. The next day, the cultures were spun down by centrifugation for 10 minutes at 4000 rpm to pellets. Ethanol (70% v/v) was added and the pellets were re-suspended and vortexed to ensure complete immersion. 4 hours later, the pellets were spun down, ethanol was discarded and the pellet was washed by immersing in Phosphate-buffered Saline (PBS), spinning down to obtain a pellet and discarding the PBS. The wash step was repeated and the final pellet was re-suspended in 100mg/ml solution using PBS, serially diluted and plated on YCFA media in anaerobic conditions supplemented with sodium taurocholate to stimulate spore germination. Ethanol resistance was determined by counting colonies (indicating germinated spores) that were present the following day. To account for species that don’t sporulate *in vitro*, if a species was originally cultured from ethanol treated faeces it was considered spore-forming. Species that didn’t survive ethanol shock treatment were also checked to see if they were originally cultured from non-ethanol treated faeces only.

### TEM

Spore images were generated using transmission electron microscopy (TEM) as previously described^63^. Bacterial isolates were streaked from frozen glycerol stocks on YCFA media^32^ in anaerobic conditions in an A95 Whitley Workstation and purity was confirmed by morphological examination and full length 16S ribosomal RNA gene sequencing. The isolates were then inoculated in YCFA broth for 2 weeks in order to induce stress conditions and stimulate sporulation before TEM images were prepared. Genome accessions of the 6 isolates are: No.1 = ERR1022323, No.2= ERR1022375, No. 3= ERR171272, No. 4= ERR1022380, No.5= ERR1022472 and No. 6 = ERR1022333.

### Functional enrichment

To identify protein domains in a genome, RPS-BLAST using COG database^64^ (accessed April 2019) was utilised. All protein domains were classified in different functional categories using the COG database. Domain and functional enrichment analysis was calculated using one-sided Fisher’s exact test with *P* value adjusted by Hochberg method in R v. 3.2.2. Functional classes were manually assigned to genes using the functional scheme originally developed for *Escherichia coli*^65^. In total, 83% of enriched FSF genes (225/272) were assigned to classes of a known function compared to 92% of enriched SF genes (450/489).

### Paralog analysis

To identify paralogs in a genome, protein domains were identified using COG database^64^ (accessed April 2019). Paralogs were called if multiple copies of a protein domain are present in a genome. Percentage of paralogs was calculated using number of paralogs and total number of protein domains present in a genome.

### CAZyme analysis

The presence of Carbohydrate-Active Enzymes^66^ was determined by querying the dbCAN families in the HMM database using hmmscan against the amino acid sequences of the protein coding genes in the genomes. Hits were filtered based on an alignment of >80 amino acids using E-value of less than 1e-05 or E-values of less than 1e-03 covering greater than 30% of the HMM hit. dbCAN families not directly related to carbohydrate utilization were removed prior to analysis, these were all auxiliary activities, all glycosyltransferases and carbohydrate esterase 10 (CE10). This left 219 entries in total to query.

### Biolog analysis

The *Erysipelotrichaceae* isolates used for Biolog analysis are described in **Supplementary Table 3**. *Longicatena caecimuris* and *Erysipelatoclostridium ramosum* have been deposited with the NCIMB culture collection under the accessions NCIMB 15236 and NCIMB 15237 respectively as part of this study. All other isolates have been previously isolated by us in the Host-Microbiota Interactions Laboratory and deposited in public culture collections except for *Faecalitalea cylindroides* (DSM3983), *Holdemanella biformis* (DSM3989) and *Eggerthia cateniformis* (DSM20559) which were obtained from DSMZ^67,68^. The FSF selected have a sporulation signature score ranging from 19%-23%, the SF selected have a sporulation signature score ranging from 47% to 56%. Before Biolog experiments, isolates were tested for ethanol resistance or were assessed if originally isolated from ethanol treated faeces. All 4 FSF isolates did not survive ethanol exposure and were not isolated from ethanol treated faeces. For the SF, *Clostridium innocuum* was successfully isolated following ethanol exposure and *Longicatena caecimuris* was not isolated following ethanol exposure but was originally isolated from ethanol treated faeces^23^. *Clostridium spiroforme* and *Erysipeloclostridium ramosum* did not survive ethanol exposure and were not originally isolated from ethanol treated faeces by us. However, there are numerous reports of these two species forming spores in the literature including imaging^69-71^. Based on this and combined with our genomic predictions they were characterised as spore-forming.

Isolates were re-streaked on YCFA agar media and grown overnight before using (*Holdemania filiformis* was allowed to grow for 2 days until sufficient growth had occurred). Cotton swabs were used to remove colonies which were then inoculated in AN-IF Inoculating Fluid (Technopath product code 72007) to a turbidity of 65% using a turbidimeter. Then 100ul was pipetted into each well of Anaerobe AN Microplates (Technopath, product code 1007) which contains 95 different carbon sources. The plates were sealed in PM Gas Bags (Technopath, product number 3032) and run on the Omnilog system for 24 hours. For each isolate, 3-5 replicates run on different days from different starting colonies were used. Data was analysed using the CarboLogR application^72^.

### Metagenomic abundance and prevalence

For estimation of prevalence and abundance in metagenomes we used SF and FSF genomes along with a genome collection representative of the entire intestinal microbiota (which also includes non-Firmicutes such as Proteobacteria, Bacteroidetes and Actinobacteria)^45^. We first determined genome quality using CheckM^57^ (‘lineage_wf’ function) and then de-replicated at an estimated species-level^73,74^ using dRep v2.2.4^75^. Briefly, genomes with a Mash^76^ distance < 0.1 were first grouped (option ‘-pa 0.1’) and subsequently clustered at an average nucleotide identity of 95% with a minimum alignment fraction of 60% (options ‘-sa 0.95 -nc 0.60’). The best quality representative genome was selected from each cluster on the basis of the CheckM completeness, contamination and the assembly N50. Each species representative was taxonomically classified with the Genome Taxonomy Database^52^ toolkit v0.2.1 using the ‘classify_wf’ function and default parameters. Sporulation capability of the resulting 562 species-level genomes were calculated as described above.

To quantify the prevalence and abundance of each species, we aligned the sequencing reads from 28,060 metagenomic datasets to our set of 562 species representatives with BWA v0.7.16a-r1181^77^. The reference database used was first indexed with ‘bwa index’ and metagenomic reads were subsequently aligned with ‘bwa mem’. Prevalence was determined as previously described^78^, where species presence was inferred when a genome was covered across at least 60% length, allowing a maximum level of depth variation according to the percentage of the genome covered (taken as the 99^th^ percentile across all data points). Coverage and depth were inferred with samtools v1.5 and the function ‘depth’^79^.

Abundance was quantified by first filtering for uniquely mapped and correctly paired reads (‘samtools view -f 2 -q 1’) and normalized both by sample sequencing depth and genome length into Reads Per Kilobase Million (RPKM) using the following formula:

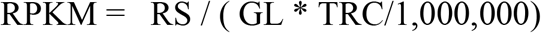

RS represents the number of reads uniquely mapped, GL the reference genome length in kilobases (kb) and TRC the total read count of the metagenomic dataset used for mapping. The level of species prevalence and abundance was compared between SF, FSF and non-spore forming non-Firmicutes bacteria using a two-tailed Wilcoxon rank-sum test. For estimating differences in abundance, only those species present in more than 10 metagenomic datasets were considered.

## Acknowledgments

This work was supported by Wellcome Trust core funding [098051] and the Australian National Health and Medical Research Council [1091097, 1159239 and 1156333 (S.C.F.) and the Victorian Government’s Operational Infrastructure Support Program (S.C.F.). The authors would also like to acknowledge the support of the Wellcome Sanger Institute Pathogen Informatics Team. We thank A. Neville for critical feedback of the manuscript.

## Funding for open access charge

Wellcome Sanger Institute.

## Author Contributions

H.P.B. and T.D.L. conceived the study. H.P.B., N.K and S.C.F. performed bioinformatics analysis. A.A. performed metagenomics analysis. K.V. analysed Biolog data. H.P.B, A.T.A., E.V. and N.J.R.D. performed *in vitro* experiments. C.C. and D.G generated TEMs. H.P.B. and T.D.L. wrote the paper with input from all authors.

